# Retention of a single *Cenp-C* gene in different syntenic locations in the *montium* group of *Drosophila* species

**DOI:** 10.64898/2025.12.11.693522

**Authors:** Rafaella F. Soares, Ching-Ho Chang, Leonardo B. Koerich, Harmit S. Malik, Gustavo C.S. Kuhn

**Author notes:** Corresponding author: Gustavo Kuhn, Universidade Federal de Minas Gerais, Av. Antônio Carlos 6627, CEP 31270-901, Belo Horizonte – MG – Brazil.

## Abstract

Chromosome segregation in eukaryotes requires the orchestrated interaction of chromosomes with microtubules, mediated by the kinetochore multiprotein complex that assembles on specific chromosomal regions known as centromeres. In most eukaryotes, two centromeric proteins, CenH3 and Cenp-C, are essential for centromere function. In *Drosophila*, the localization of CenH3 (referred to as Cid in *Drosophila*) depends on its chaperone CAL1 and Cenp-C. Previous studies have shown that both *Cid* and *Cenp-C* underwent a coincident gene duplication and likely functional specialization in the *Drosophila* subgenus. Independently, *Cid* duplications led to *Cid1, Cid3,* and *Cid4* paralogs in the *montium* group (*Sophophora* subgenus), but it is unknown whether this group also underwent parallel duplications of *Cenp-C*. Here, we investigate this possibility by analyzing sequenced genomes of 23 *montium* group species. We identified *Cenp-C* genes in five distinct syntenic loci; we named these genes *Cenp-C1b*, *Cenp-C1c*, *Cenp-C1d*, *Cenp-C1e* and *Cenp-C3*. Despite their distinct synteny, most *montium* group species only encode a single *Cenp-C*; their phylogeny mirrors the species phylogeny, and they appear to have retained Cenp-C protein motifs indicative of function. A closer examination revealed that these *Cenp-C* genes resulted from gene translocations or alternate retention (duplication followed by loss of the ancestral copy); only one species, *D. vulcana,* retains two intact *Cenp-C* paralogs. Therefore, unlike the *Drosophila* genus, the co-retention of three *Cid* paralogs in the *montium* group has not resulted in a coincident *Cenp-C* paralog co-retention. Our work highlights differences in functional retention and likely specialization of the two most conserved centromeric proteins in eukaryotes.

## Introduction

High-fidelity chromosome segregation during cell division is crucial for life. In eukaryotes, this process relies on centromeres – chromosomal regions that orchestrate kinetochore assembly and attachment to spindle microtubules to ensure accurate chromosome segregation. The essential function of the centromere contrasts with the moldable structure that defines it across the vast diversity of taxa (Akiyoshi & Gull, 2014; Drinnenberg et al., 2014). Despite some prominent exceptions, centromeric DNA in most eukaryotes is wrapped by a centromere-specific histone H3 (CenH3) variant, Cenp-A (Earnshaw & Rothfield, 1985; Palmer et al., 1987), which replaces histone H3 in centromeric chromatin. Indeed, the establishment of a functional centromere is determined primarily by the presence of CenH3 rather than by underlying sequences in most eukaryotes (Rosin & Mellone, 2017). CenH3 is also critical for the subsequent localization of other kinetochore components (McKinley & Cheeseman, 2016) and is therefore indispensable for centromeric function in all species that encode CenH3.

Duplications of *CenH3* have been reported in some plants (Finseth et al., 2015; Neumann et al., 2015) and animals (Caro et al., 2022; Kursel et al., 2020; Kursel & Malik, 2017; Monen et al., 2015; Teixeira et al., 2018). In *Drosophila* species, *CenH3* (or *Cid*, for *centromere identifier;* Henikoff et al., 2000) is present either as a single-copy gene, as in *D. melanogaster*, or in multiple copies (Kursel & Malik, 2017; Teixeira et al., 2018). For example, five independent duplications of *Cid1* (the ancestral copy) led to the emergence of *Cid2*, *Cid3*, *Cid4, Cid5,* and *Cid6* in different *Drosophila* species. *D. eugracilis* encodes *Cid2*, but its ancestral *Cid1* has undergone pseudogenization; thus, it only encodes a single functional *Cid* gene. Species from the *Drosophila* subgenus encode two functional copies (*Cid1* or *Cid6,* and *Cid5*), while species from the *montium* group present three functional copies (*Cid1*, *Cid3,* and *Cid4*) (Kursel & Malik, 2017; Teixeira et al., 2018). In some species, *Cid* paralogs experienced substantial divergence and now perform non-redundant centromeric roles. For example, in *D. virilis* from the *Drosophila* subgenus, *Cid1* performs a centromeric function in somatic cells and throughout female gametogenesis, whereas *Cid5* performs a centromeric function throughout male gametogenesis, including the inheritance of centromeric identity through sperm (Kursel et al., 2021; Kursel & Malik, 2017). Although not as well studied as in the *Drosophila* subgenus, specialized expression of *Cid* paralogs (*Cid1, Cid3,* and *Cid4*) in the *montium* group also suggests their functional specialization (Kursel & Malik, 2017).

Another essential centromeric protein found in most eukaryotes is Cenp-C, whose interaction with both CenH3 and centromeric DNA seems to be necessary to mediate Cenp-C association with centromere chromatin (Kato et al., 2013; Medina-Pritchard et al., 2020; Politi et al., 2002; Roure et al., 2019). Studies in insects have revealed the strong interdependence between Cenp-C and CenH3 for their centromeric function. For example, Cenp-C is crucial for targeting and deposition of CenH3 onto an existing centromere in *D. melanogaster* (Erhardt et al., 2008; Roure et al., 2019). Conversely, Cenp-C localization to the centromere also depends on CenH3 (Erhardt et al., 2008; Roure et al., 2019).

Species from the *Drosophila* subgenus not only encode two copies of *Cid* (*Cid1* and *Cid5/Cid6*), but also two copies of *Cenp-C* (*Cenp-C1* and *Cenp-C2*) (Teixeira et al., 2018) that appear to have co-diverged and co-specialized across phylogeny. Although these two *Cenp-C* paralogs are highly divergent, they retain key motifs that might play important roles in centromere localization and function. Both paralogs are expressed in almost all developmental stages. However, *Cenp-C2* expression is male-biased, raising the possibility that Cenp-C2 interacts with Cid5 during male gametogenesis, whereas Cenp-C1 likely retains its canonical Cenp-C function (Teixeira et al. 2018). In this scenario, the co-retention of Cid and Cenp-C duplicates might be related to their functional interactions and specializations. A functional co-dependence between CenH3 and Cenp-C is further reinforced by studies showing a coincident loss of both CenH3 and Cenp-C in insect species that have transitioned from monocentric to holocentric chromosomes (Drinnenberg et al., 2014). However, co-duplication or co-retention of CenH3 and Cenp-C is not observed in all cases. For example, many plant species encode duplicates of one but not the other protein (Ishii et al., 2020; Talbert et al., 2004). Similarly, despite recurrent duplication of both CenH3 and Cenp-C in *Caenorhabditis* nematodes, duplication or retention of these two genes does not appear to be strongly correlated (Caro et al., 2022).

Nevertheless, the strong interdependence between CenH3 and Cenp-C, and the co-divergence of their paralogs in the *Drosophila* subgenus, prompted us to investigate whether a similar co-divergence and functional specialization may have occurred in the *montium* group of the *Sophophora* subgenus in *Drosophila*, where three *Cid* genes have been co-retained (Kursel & Malik, 2017). With 94 described species, the *montium* group is the largest clade in the *Sophophora* subgenus. It is estimated to have diverged from the *melanogaster* group around 28 million years ago (Russo et al. 2013) and is subdivided into seven subgroups: *montium*, *parvula*, *kikkawai*, *serrata*, *punjabiensis*, *seguyi,* and *orosa* (Yassin, 2018). The monophyly of the *montium* group is strongly supported (Conner et al., 2021; Da Lage et al., 2007; Finet et al., 2021; Russo et al., 2013; Yang et al., 2012). A recent study analyzing 60 genes also supports the monophyly of all seven subgroups proposed by Yassin (2018), except for one species (*D. baimaii*) out of 42 (Conner et al., 2021). The recent sequencing of genomes from 23 species in the *montium* group, including representatives of almost all *montium* subgroups (Bronski et al., 2020), prompted us to conduct an in-depth analysis of the duplication and retention of *Cenp-C* paralogs in these species.

Our analyses initially revealed apparent *Cenp-C* duplications within the *montium* group, with *Cenp-C* genes located in distinct syntenic regions of the genome across various species. However, closer examination revealed a more complex scenario: most species in the *montium* group encode only one functional *Cenp-C* gene, which encodes most of the protein motifs required for function and whose phylogeny is consistent with the proposed species’ phylogeny. The only exception was *D. vulcana*, which encodes two Cenp-C paralogs. We conclude that repeated relocation of *Cenp-C* to new genomic loci occurred either by genetic translocations or by alternate retention, with new copies of *Cenp-C* in new genomic locations being retained while the ancestral *Cenp-C* gene was lost. Thus, despite substantial opportunities to co-diversify with *Cid* genes in the *montium* group, *Cenp-C* paralogs have failed to do so. Our findings reiterate differences in the hierarchy of functional specialization between Cid and Cenp-C. They also highlight how different mutations can lead to co-retention and functional specialization of paralogs in some instances, but not in others.

## Results

### *Cenp-C* genes are present in distinct genomic locations in the *montium* group of *Drosophila* species

A previous study analyzed 17 species from the *montium* group to reveal that these species possess three functional copies of *Cid* (*Cid1*, *Cid3*, and *Cid4*) (Kursel & Malik, 2017). We investigated the retention of *Cid* paralogs in an additional 10 species that were not previously analyzed but are included in the 23 *montium* group sequenced species studied here. Using *tBLASTn* searches in combination with previously identified syntenic regions for *Cid1*, *Cid3*, and *Cid4* paralogs, we found that 9 of the 10 species contain *Cid1*, *Cid3*, and *Cid4* in their genomes. The only exception was *D. bunnanda*, which encoded intact *Cid1* and *Cid4*, but a *Cid3* pseudogene **(Supplementary Figure 1)**. This is not unexpected as *D. bunnanda*’s sister species, *D. mayri,* was also previously shown to encode a *Cid3* pseudogene (Kursel & Malik, 2017). Except for *D. bunnanda* and *D. mayri*, we infer that all *montium* group species encode three functional copies of *Cid*.

Given the previously documented co-retention and potential co-specialization of *Cid* and *Cenp-C* paralogs in species from the *Drosophila* subgenus (Teixeira et al. 2018), we investigated whether *montium* group species also underwent a parallel co-retention of *Cenp-C* paralogs. As a first step to revealing putative *Cenp-C* paralogs in the *montium* group of species, we first identified the shared syntenic location of *Cenp-C* genes outside the *montium* group in four species: two from the *Sophophora* subgenus (*D. melanogaster* and *D. ananassae*), one from the *Drosophila* subgenus (*D. virilis*), and a basal species (*Scaptodrosophila lebanonensis*). Their *Cenp-C* genes (also called *Cenp-C1* in *D. virilis*) are flanked on the 3’ end by the *5-HT2B* gene and on the 5’ end by *CG1427* in three of these four species. In *D. melanogaster,* it is flanked on the 5’ end by *CG31640* because of a subsequent lineage-specific chromosomal alteration. Based on this, we conclude that the ancestral syntenic location of *Cenp-C1* in *Drosophila* species was between the *CG1427* and *5-HT2B* genes in the Muller element E, corresponding to chromosome arm 3R in *D. melanogaster* (we refer to *Cenp-C1* found at this location as *Cenp-C1a*) (**Figure 1A**). Species from the *Drosophila* subgenus also encode a *Cenp-C2* paralog located between the *CLS* and *RpL27* genes in *D. virilis,* also on Muller element E (Teixeira et al., 2018) **(Figure 1A)**.

**Figure 1.**
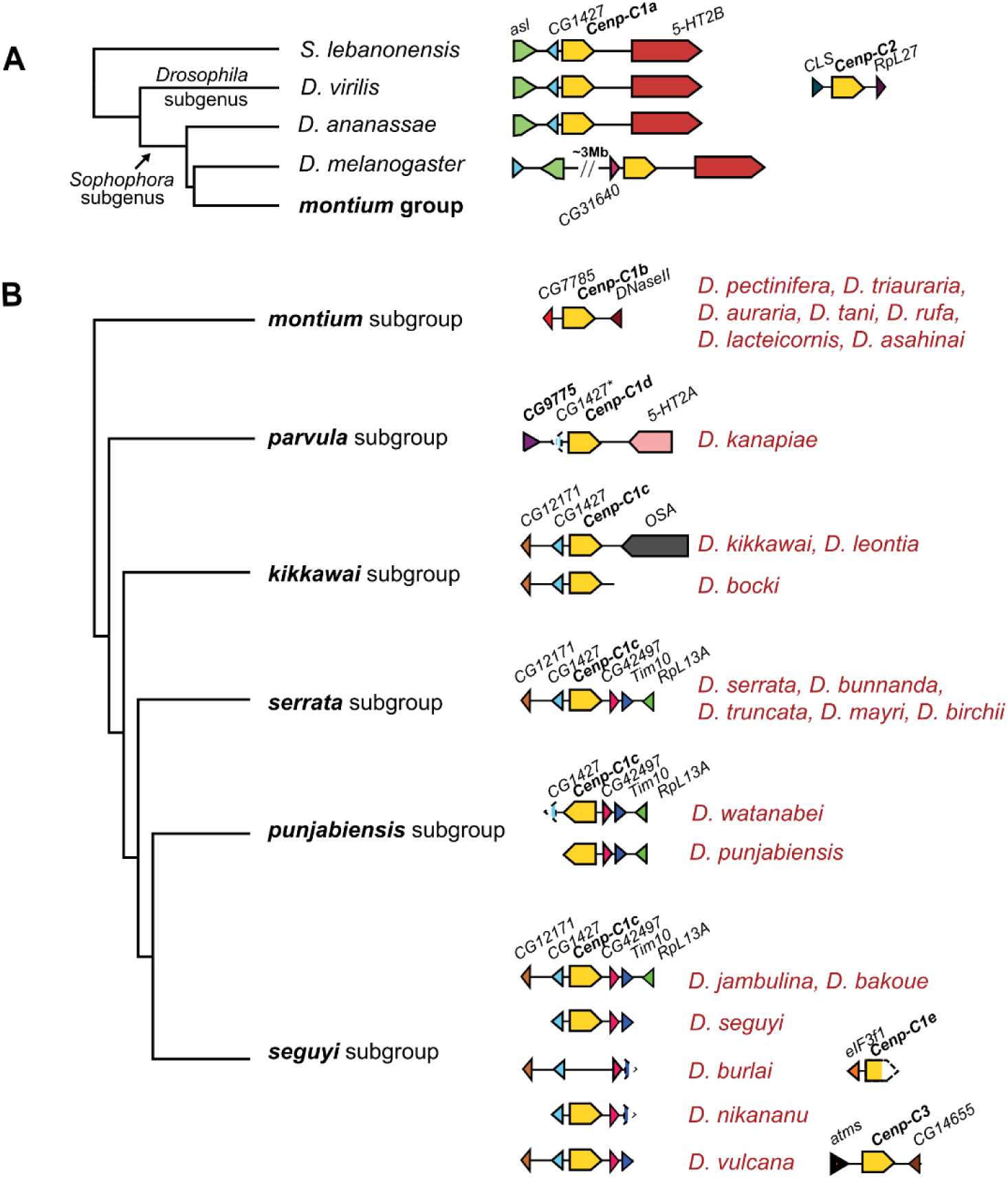
Location of *Cenp-C* genes in *Drosophila* species. **(A)** Schematic representation of the *Cenp-C* locus in four outgroup species of the montium group of *Drosophila* species. **(B)** Phylogeny of the *montium* subgroups (Conner et al. 2021), with a schematic representation of the genomic loci where *Cenp-C* genes have been identified in the present study. Genes are shown as arrows. Potential pseudogenes are indicated with an asterisk (*e.g., CG1427* in *D. kanapiae*) while incomplete genes due to missing contigs are indicated with dashed boxes (*e.g., Cenp-C1e* in *D. burlai*).

The *melanogaster, montium,* and *ananassae* groups of *Drosophila* species form a near-trichotomy, with the *ananassae* group as an outgroup to the other two groups (Kopp 2006). Based on this, we expected to find *Cenp-C* orthologs at the same syntenic location as *D. ananassae*. To test this expectation, we identified *Cenp-C* in 24 species across six of the seven subgroups in the *montium* group (all but *orosa*): *montium*, *serrata*, *punjabiensis*, *seguyi*, *kikkawai*, and *parvula* (Yassin, 2018). In the *montium* subgroup, we identified a single *Cenp-C* gene (tentatively renamed *Cenp-C1b*), but in a new genomic location, flanked by the *CG7785* and *DNaseII* genes **(Figure 1B)**. We could also assemble a single *Cenp-C* locus from three contigs in *D. kanapiae* from the *parvula* subgroup **(Supplementary Figure 2A).** Based on this reconstruction, we concluded that *Cenp-C* is flanked by a fragment of *CG1427* at the 5’ end as well as *5-HT2A* at its 3’ end. Since the *CG1427* is not intact and this shared syntenic location is distinct from any of the other *Cenp-C* genes described earlier, we named this copy *Cenp-C1d* to distinguish it from other *Cenp-C* locations **(Figure 1B)**.

*Cenp-C* genes in the *serrata*, *kikkawai*, *punjabiensis*, and *seguyi* subgroups appear to map to similar syntenic locations. In the *serrata* subgroup, the single *Cenp-C* gene is flanked by the *CG1427* gene at its 5’ end, like the ancestral *Cenp-C1a* locus, but by *CG42497, Tim10,* and *RpL13A* genes (instead of *5-HT2B*) at its 3’ end **(Figure 1B)**. To be conservative, we refer to this gene in the *serrata* subgroup as *Cenp-C1c* to distinguish it from *Cenp-C1a, Cenp-C1b,* and *Cenp-C1d*. Similarly, a single *Cenp-C* gene is present in species from the *D. kikkawai* subgroup, flanked on the 5’ end by the same genes as *Cenp-C1c* in the *serrata* subgroup, even though it is flanked by the distinct *OSA* gene at its 3’ end **(Figure 1B)**. Despite the short contig sizes, we could confirm that the *punjabiensis* subgroup species share the same *Cenp-C1c* gene and location as the *serrata* subgroup, except that the orientation of the *Cenp-C1c* gene is reversed **(Figure 1B)**. The *Cenp-C1c* and *Cenp-C1d* syntenic locations share *CG1427* with the ancestral *Cenp-C1a* syntenic location (outside the *montium* group). However, in the absence of additional syntenic markers, we cannot infer that *Cenp-C1a, Cenp-C1c,* and *Cenp-C1d* are orthologous based on this evidence alone. For example, it is just as likely that *Cenp-C* and *CG1427* co-duplicated or relocated to a new syntenic location; we discuss these possibilities later in this report.

We readily identified a single copy of *Cenp-C1c* in most *seguyi* subgroup species **(Figure 1B**). However, in *D. burlai from the seguyi subgroup*, we were unable to identify the *Cenp-C1c* gene in its syntenic location. Since *Cenp-C* is essential in *Drosophila*, this suggests that *D. burlai* might encode its *Cenp-C* gene in a different syntenic location. Indeed, we did identify a 3,322 bp contig encoding the 5′ fragment of *Cenp-C,* with no obvious mutations, flanked on the 5’ end by *eIF3f1* **(Figure 1B)**. Since this fragment resides at a distinct genomic location from *Cenp-C1c,* it may represent yet another syntenic region for *Cenp-C* in *D. burlai*, which we tentatively name *Cenp-C1e* **(Figure 1B)**. However, in the absence of complete sequence data for the full-length *Cenp-C* gene, we do not consider this gene further in our study. The *D. burlai* case suggests that the diversity of syntenic loci in which we have uncovered *Cenp-C* genes in the *montium* group of species may still be an underestimate.

Based on these findings, we conclude that, unlike other *Drosophila* species, the *Cenp-C* locus has recurrently undergone changes in syntenic location within the *montium* group. The early branching *montium* and *parvula* subgroups encode *Cenp-C1b* and *Cenp-C1d* in distinct locations, whereas the common ancestor of the *serrata, kikkawai, punjabiensis,* and *seguyi* subgroups all encoded a single *Cenp-C1c* gene. All these syntenic locations map to Muller element E (https://flybase.org/jbrowse). With one or a few exceptions (discussed below), all *montium* group species appear to encode a single intact *Cenp-C* gene, which contrasts with their retention of two or three intact *Cid* genes.

### Co-retention of multiple *Cenp-C* paralogs within *montium* group species

Our survey revealed only one species within the *montium* group – *D. vulcana* from the *seguyi* subgroup – with robust evidence of co-retention of two *Cenp-C* paralogs. In addition to the *Cenp-C1c* gene, also found in other members of the *seguyi* subgroup, we found that *D. vulcana* encodes a second copy of *Cenp-C* (which we refer to as *Cenp-C3*), split across two contigs. The breakpoint in the sequence corresponds to a predicted intron sequence in the *Cenp-C3* gene (**Supplementary Figure 2B**). Although we could not fully reconstruct the predicted intron sequence due to the absence of an overlapping sequence, both ends of *Cenp-C3* are flanked by annotated genes, at the 5’ end by *atms* and at the 3’ end by *CG14655. CG14655* and *atms* are neighboring genes in other *montium* group species. Based on this evidence, we conclude that *D. vulcana* is exceptional in encoding both *Cenp-C1c,* like other members of the *seguyi* subgroup, and *Cenp-C3* in a new syntenic location flanked by *atms* and *CG14655* **(Figure 1B)**.

Both *Cenp-C1c and Cenp-C3* genes in *D. vulcana* appear to be intact. We found no evidence for frameshifts or stop codons that would disrupt the open reading frame of either *Cenp-C1c* or *Cenp-C3*. However, our analysis using the gene prediction algorithm (Augustus) initially failed to detect three exons in *Cenp-C1c,* which are present in other *Cenp-C1c* genes and in *Cenp-C3* **(Supplementary Figure 3)**. Additionally, these three exons are found in the opposite orientation within a predicted intron of the *Cenp-C1c* gene (**Supplementary Figure 2C**), flanked by similar transposable elements (TEs) on both sides. Based on this evidence and the conservation of the three exons in other *Cenp-1c* orthologs, we conclude that this sequence segment containing the three exons was incorrectly inverted during the assembly of the *D. vulcana* sequenced genome and that *Cenp-C1c* and *Cenp-C3* likely encode the same number of exons. Alignment of the amino acid sequences encoded by *Cenp-C1c* and *Cenp-C3* from *D. vulcana* revealed nearly 90% amino acid identity. Moreover, protein motif analysis (see below) revealed the preservation of six of the motifs associated with essential centromeric function in *D. vulcana* Cenp-C1c and Cenp-C3. Therefore, preliminary evidence suggests that both *Cenp-C* paralogs in *D. vulcana* are functional.

*Cenp-C3* appears to be the only *bona fide Cenp-C* paralog in the *montium* group of *Drosophila* species. Our genomic search for *Cenp-C* paralogs retrieved several other contigs containing small fragments of *Cenp-C* in four species: *D. burlai, D. punjabiensis*, *D. vulcana*, and *D. bakoue* **(Supplementary Figure 4)**. In *D. burlai*, we identified two short contigs containing *Cenp-C* fragments that lack most of the 5′ portion of the gene and are immediately flanked by transposable elements and different neighboring genes, suggesting that these fragments may be pseudogenes. In *D. punjabiensis* (*punjabiensis* subgroup), we identified an incomplete *Cenp-C* flanked by *Grip84* and *Rab23*, located near the *Cenp-C1c* locus. In *D. bakoue* (*seguyi* subgroup), we identified a *Cenp-C* fragment within a contig, along with the gene *canoe* (*cno*). Finally, in *D. vulcana* (*seguyi* subgroup), we found a *Cenp-C* fragment flanked on one side by eight additional genes (*Sec63*, *Sh3β, hzg*, *Hip14*, *PolD1*, *Arl1*, *CG17027*, and *CG17029* located on the Muller element D). None of these contigs appears to encode intact *Cenp-C* paralogs. Still, they confirm the success of our homology-based approaches to identify even small segments of *Cenp-C* pseudogenes in the *montium* group species.

### Conservation of Cenp-C functional motifs

We next assessed whether all the various *Cenp-C* genes we have identified in different syntenic locations are likely to be functional. First, we confirmed that none of them have any mutations that would compromise their open reading frame. Next, we compared protein motif conservation across the proteins encoded by these genes. *Drosophila* Cenp-C proteins include seven conserved motifs (Heeger *et al*. 2005): arginine-rich (R-rich), *Drosophilid* Cenp-C homology (DH), two predicted AT-hook domains (AT1 and AT2), Nuclear Localization Signal (NLS), Cenp-C motif (essential for CenH3 binding), and a dimerization domain near the C-terminal region (Cupin or C-term) (**Figure 2A**). The Cupin motif is essential for Cenp-C interaction with the Cal1 chaperone for the deposition of Cid nucleosomes at the centromere (Medina-Pritchard et al., 2020; Roure et al., 2019). The specific functions of the remaining motifs (R-rich, DH, NLS, AT1, and AT2) have yet to be elucidated. However, Cenp-C variants lacking any of these motifs (except AT1) exhibit phenotypic abnormalities in mutant embryos (Heeger et al., 2005).

**Figure 2.**
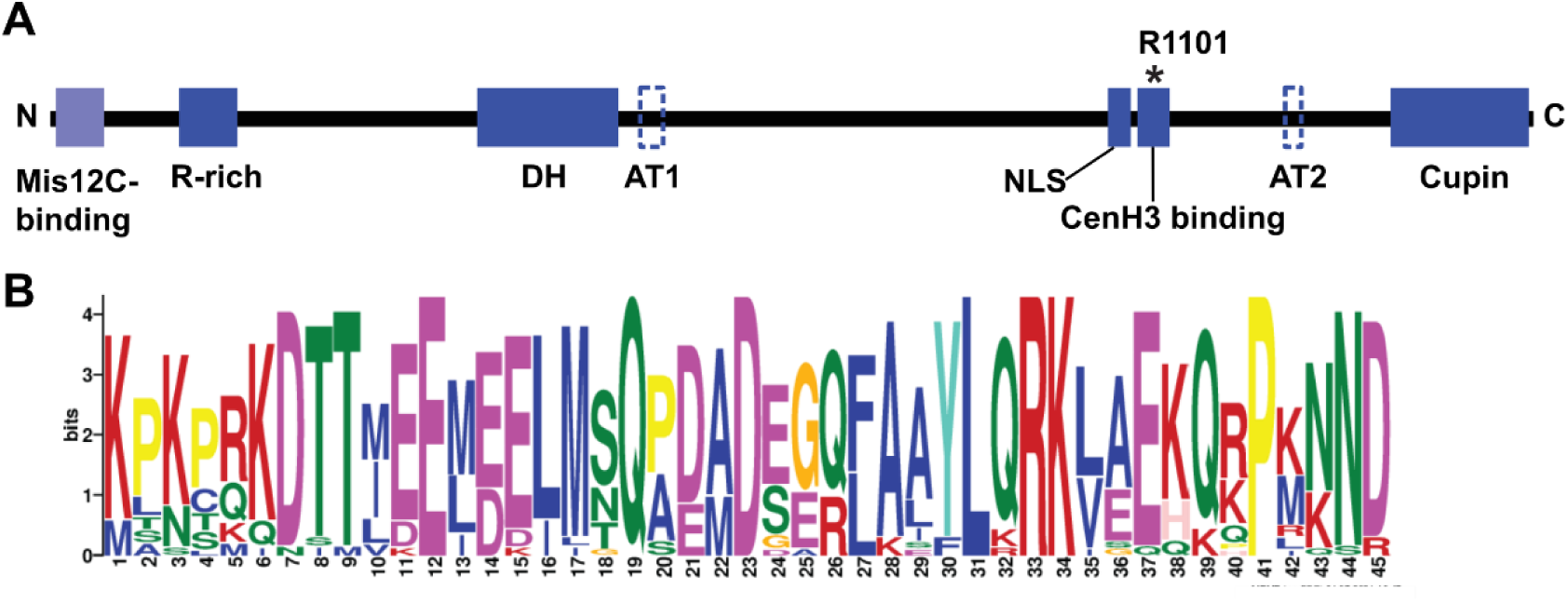
*Drosophila* Cenp-C functional motifs. **(A)** Representation of the seven functional motifs in Cenp-C identified in *D. melanogaster* (Heeger et al. 2005), along with an additional N-terminal Mis12C-binding motif identified in the present study. AT1 and AT2 motifs represent AT-hook domains that could not be identified in species from the *montium* group. The black asterisk marks arginine (R1101) in Cenp-C, which is identical across all Cenp-C orthologs and paralogs. N and C denote the N-terminal and C-terminal regions of the protein, respectively. **(B)** Sequence Logo generated by MEME for the putative Mis12-binding motif (Mis12C binding).

We found that all these motifs, except AT1 and AT2, are conserved in the Cenp-C genes from the *montium* group species. In particular, the Cenp-C motif, including a critical R1101 residue required for Cenp-C localization to the centromere (Heeger et al., 2005), is conserved (**Figure 2A, Supplementary Figure 5**). Since the AT1 and AT2 motifs are predicted to bind AT-rich DNA in the minor groove (Heeger et al., 2005), their activity might be replaced by other non-specific DNA-binding motifs. In addition to the previously defined motifs, we also detected a new motif in the N-terminal region of all Cenp-C proteins from the *montium* group. This motif corresponds to the first 45 amino acids from the *Drosophila melanogaster* Cenp-C1a (**Figure 2B, Supplementary Figure 5**), which were previously shown to be important for recruiting core kinetochore proteins to the centromere by directly binding to Mis12C (Liu et al., 2016; Przewloka et al., 2011). These findings were corroborated by another study, which demonstrated that the first 45 amino acids of *D. melanogaster* Cenp-C1a interact with Mis12-Nnf1a (Mis12C subunits) (Richter et al., 2016). Therefore, the conservation of this N-terminal motif in Cenp-C is consistent with its predicted functional role, even though it was previously not identified in evolutionary motif analyses (**Figure 2, Supplementary Figure 5**).

### *Cenp-C* phylogeny mirrors the phylogeny of *montium* group species

Thus far, our analyses have revealed single *Cenp-C* genes in distinct syntenic locations. These genes could result from orthologs being relocated to different genomic locations or from lineage sorting and alternate retention of paralogs in different species. To help differentiate between these possibilities, we compared the phylogeny of the various *Cenp-C* genes (*Cenp-C1b*, *Cenp-C1c*, *Cenp-C1d,* and *Cenp-C3*) with that of the *montium* group species from which they were obtained. If each of these genes represents a true ortholog, we would expect their phylogeny to closely mirror the species phylogeny.

We performed a phylogenetic analysis using maximum likelihood (ML) of all *Cenp-C* paralogs from the *montium* group species, together with *Cenp-C* from all other *Drosophila* species previously studied (Teixeira et al., 2018) (**Figure 3**). In agreement with the orthology hypothesis, we found that the *Cenp-C* copies from the *montium* group form a monophyletic group, sister to the *Cenp-C1a* of all other species from the *Sophophora* subgenus. Moreover, the ML tree revealed that the different copies of *Cenp-C* form monophyletic clades in accordance with the previously proposed *montium* subgroups (Yassin 2018) and the proposed phylogeny for species of the *montium* group (Conner et al., 2021). Moreover, at least one copy of *Cenp-C* is found in the genome of each species, consistent with its essential centromere function. All protein motifs involved in centromere function, including those necessary for Cenp-C or Cid localization at the centromere, are conserved in all *Cenp-C* genes found in the *montium* group. We conclude that all *montium* group species *Cenp-C* genes (except *Cenp-C3*, which arose via a recent duplication) are functionally orthologous to *Cenp-C1a* despite being in distinct syntenic locations.

**Figure 3.**
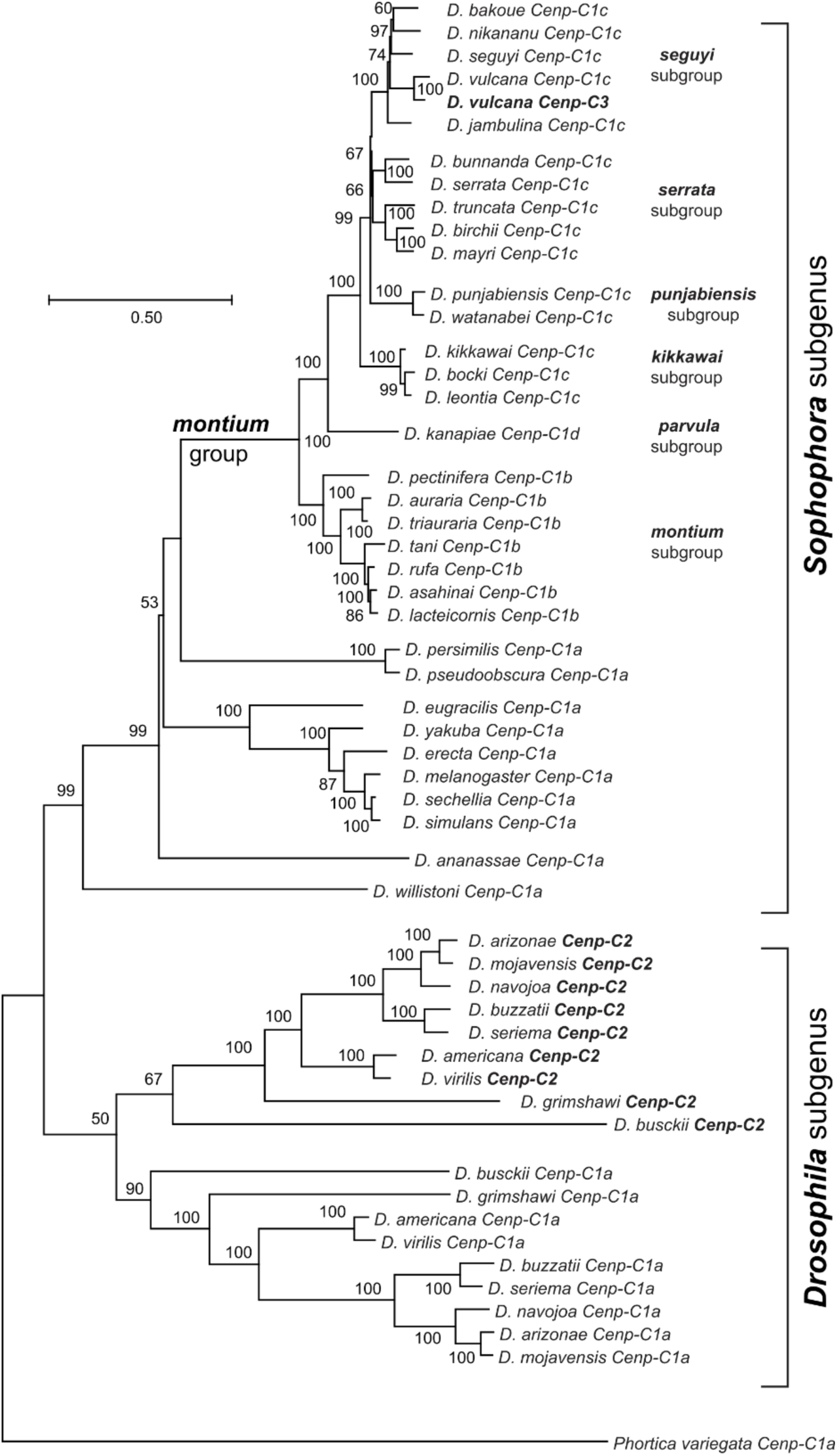
Maximum likelihood tree showing the phylogenetic relationships among *Cenp-C* sequences. The phylogeny includes *Cenp-C* genes from multiple subgroups of the *montium* group of *Drosophila* species, as well as representative species from the *Sophophora* and *Drosophila* subgenera, with *P. variegata* as an outgroup. Bootstrap values are indicated at each node except when they were below 50% (out of 1000 replicates). The scale bar represents the number of substitutions per site.

### Reconciling apparent orthology of *Cenp-C* genes despite distinct syntenic locations in the *montium* group

We wished to reconcile our observations of apparent orthology of the various *Cenp-C* genes we have identified (*Cenp-C1b*, *Cenp-C1c*, and *Cenp-C1d*) with their distinct syntenic locations. Since all *Cenp-C* genes still carry introns in homologous locations, we could rule out a hypothesis of retro-duplication via RNA intermediates. This still left two hypotheses. First, *Cenp-C* could have translocated to a new genomic location, either alone or together with neighboring genes. Alternatively, *Cenp-C* could have duplicated, with the paralog landing in a new location, followed by the loss of the ancestral *Cenp-C* gene. The only way to distinguish between these possibilities would be to find vestiges of the original gene at its original genomic location. Despite our success at finding *Cenp-C* pseudogenes using homology-based searches, the rate of pseudogene decay is rapid in *Drosophila* genomes (Harrison et al., 2003; Petrov et al., 1998; Sisu et al., 2014). Thus, the more ancient the event, the less likely we would be to find evidence for the second, alternate retention hypothesis. Nevertheless, analyses of syntenic regions where different *Cenp-C* paralogs are found could provide additional insight into distinguishing between relocation, gene duplication, and alternative retention.

To retrace the steps leading to the distinct syntenic locations of *Cenp-C* and neighboring genes, we compared the syntenic locations of all *Cenp-C* genes we have identified (*Cenp-C1b*, *Cenp-C1c*, *Cenp-C1d, and Cenp-C3*) across all *montium* group species (see **Supplementary Table 1** for scaffold IDs). Based on this, we can reconstruct the following chronology of events leading to the current status of *Cenp-C* genes in the *montium* group in a series of five steps (**Figure 4**). Based on its location in the outgroup species, we infer that the common ancestor of the *montium* group encoded a single *Cenp-C1a* gene flanked by *CG1427* and *5-HT2B* on Muller element E (**Figure 4)**. In step 1, this hypothetical ancestor then acquired the *CG42497* and *Tim10* genes from Muller element C, so that the original *Cenp-C1a* locus in the *montium* group ancestor encoded *CG1427, Cenp-C1a, CG42497, Tim10, and 5-HT2B* (**Figure 4**). This inference is based on two findings. First, *CG42497* and *Tim10* are located on Muller element C in species outside the *montium* group, including *D. melanogaster*, *D. ananassae*, *D. virilis*, and *S. lebanonensis.* Second, Muller element C in the *montium* group still encodes a copy of *Tim10* but not *CG42497* **(Figure 4)**. Therefore, we conclude that *CG42497* and *Tim10* were duplicated from the Muller element C to the *Cenp-C1a* locus in this lineage, following which *CG42497* was lost or degenerated from Muller element C (**Figure 4**).

**Figure 4.**
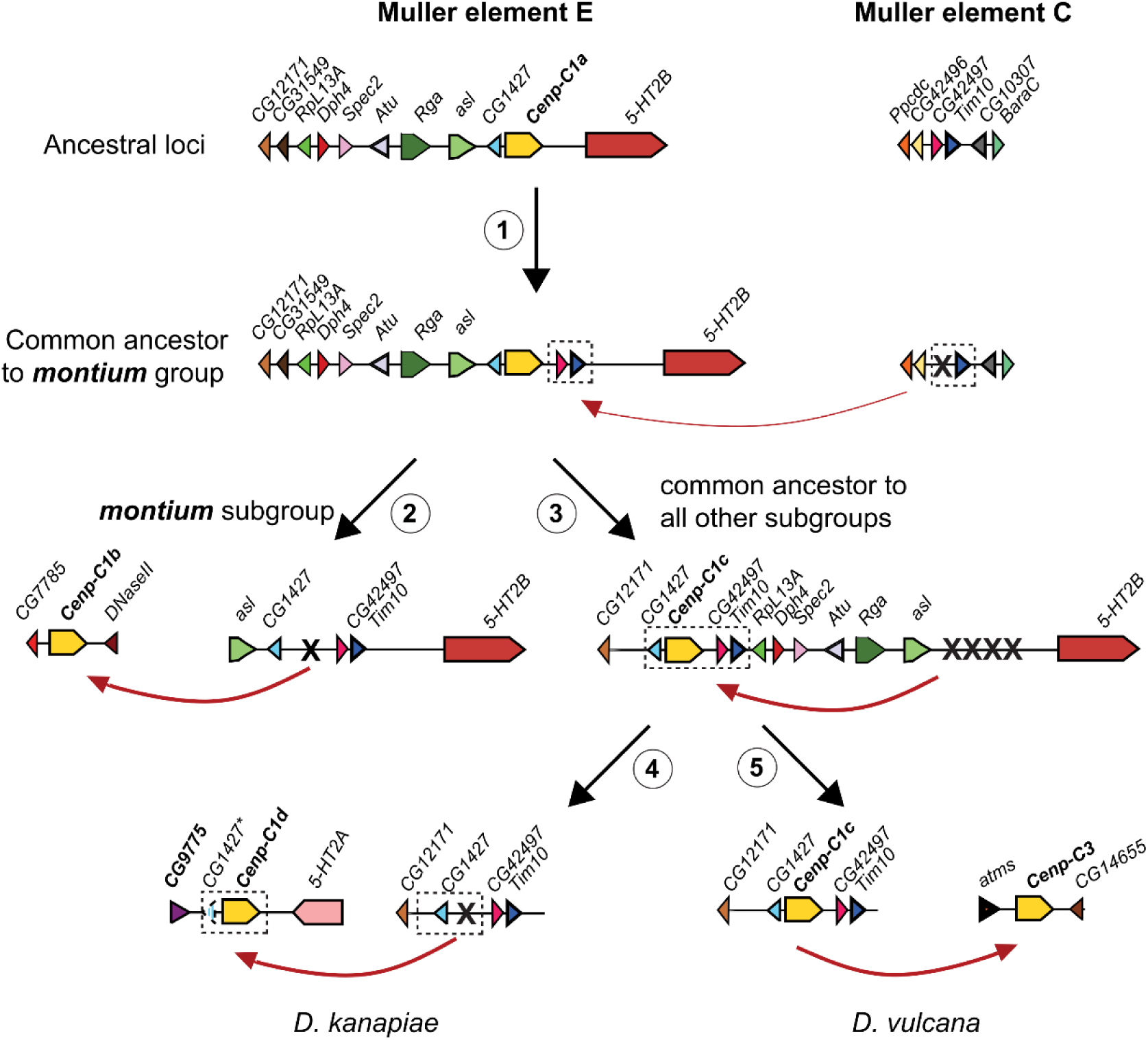
Five gene transposition or duplication events leading to distinct genomic locations of *Cenp-C* in the *montium* group. **(1)** *Cenp-C1a* is in the ancestral locus. In the common ancestor of the *montium* group, the *CG42497*/*Tim10* genes were duplicated from Muller element C to Muller element E. Subsequently, *CG42497* was lost from Muller element C. **(2)** In the *montium* subgroup, *Cenp-C1a* was either relocated or duplicated to a new locus, giving birth to *Cenp-C1b.* The *Cenp-C1a* was subsequently lost in these species. **(3)** In all other five *montium* subgroups, *Cenp-C1a* and its flanking genes (*CG1427* and *CG42497*/*Tim10*) were relocated to a new locus. All four genes were then lost from the original locus. This relocated copy of *Cenp-C* was named *Cenp-C1c*. **(4)** In *D. kanapiae,* both *Cenp-C1c* and *CG1427* were duplicated to a new locus, given rise to *Cenp-C1d*, while *Cenp-C1c* was lost in this species. The dashed blue triangle represents a *CG1427* pseudogene at the new locus (indicated with an asterisk). **(5)** In *D. vulcana, Cenp-C1c* was duplicated to a new locus, giving rise to *Cenp-C3*; both *Cenp-C1c* and *Cenp-C3* copies are maintained in *D. vulcana*. Not shown is a gene relocation or duplication of *Cenp-C1c* in *D. burlai*, giving rise to *Cenp-C1e* in a new locus (Figure 1B) followed by loss of the original *Cenp-C1c* gene.

No extant *montium* group species now encodes *Cenp-C1a* at the original location. In step 2, *Cenp-C1a* was lost from its original syntenic locus in the earliest-branching *montium* subgroup; we can find *CG1427, CG42497,* and *Tim10,* but not *Cenp-C1a*, in the original *Cenp-C1a* syntenic locus **(Figure 4)**. Instead, *Cenp-C1* is now present at a new *Cenp-C1b* locus in the *montium* subgroup, where *CG7785* and *DNaseII* flank it. Since this loss/gain involved a single gene, we cannot distinguish whether *Cenp-C1b* arose from gene relocation or from duplication/alternate retention.

Next, in step 3, we infer that most species from the *parvula, kikkawai*, *serrata*, *punjabiensis,* and *seguyi* subgroups underwent a translocation of four genes (*CG1427, Cenp-C1a, CG42497, Tim10*) from the *Cenp-C1a* to the *Cenp-C1c* syntenic locus. As a result, species from the *kikkawai*, *serrata*, *punjabiensis,* and *seguyi* subgroups now encode *CG1427, Cenp-C1c, CG42497,* and *Tim10* in the new *Cenp-C1c* location, flanked by *CG12171* and/or *RpL13A* **(Figure 4)**. Although our findings cannot formally rule out the duplication-and-alternate-retention hypothesis, we believe it is unlikely in this case, as we cannot detect any vestiges of any of these four genes at the original *Cenp-C1a* location in the *kikkawai*, *serrata*, *punjabiensis,* and *seguyi* subgroups. Under the alternate retention hypothesis, we would not necessarily expect all four genes to be co-retained in the new locus but lost in the original location.

We infer that the *Cenp-C1a* to *Cenp-C1c* translocation (step 3) was also shared with the *parvula* subgroup (including *D. kanapiae*), which is believed to be the second to branch after the *montium* subgroup), because we can find the same arrangement of *CG1427, CG42497, and Tim10* in the original *Cenp-C1c* syntenic locus. However, we no longer find *Cenp-C1c* next to *CG1427, CG42497, and Tim10* in the original *Cenp-C1c* syntenic locus. Instead, we find *CenpC1d* and a fragment of *CG1427* in a new *Cenp-C1d* location, flanked by *CG9775* and *5-HT2A* **(Figure 4)**. Here, we favor the duplication and alternate retention hypothesis, in which a duplication of *Cenp-C1c,* along with a full or partial copy of *CG1427,* occurred at the *Cenp-C1d* location in *D. kanapiae,* followed by the loss of *Cenp-C1c* in the original locus and the ongoing degeneration of *CG1427* in the ‘new’ *Cenp-C1d* locus (step 4, **Figure 4)**. In *D. burlai*, we can find *CG1427, CG42497, and Tim10,* but not *Cenp-C1c*, in the original *Cenp-C1c* syntenic locus. Therefore, we cannot rule out whether *Cenp-C1e* arose from relocation or from duplication/alternate retention in these species (**Figure 1B**).

Thus, we conclude that both single- and multi-gene relocations or duplications, followed by alternate retention, have occurred to maintain functional orthology between the *Cenp-C1b, Cenp-C1c,* and *Cenp-C1d* genes and the outgroup species’ *Cenp-C1a* genes, despite them all being found in distinct syntenic locations. In the final, most recent step (step 5, **Figure 4**), the *Cenp-C1c* gene duplicated, giving rise to the paralogous *Cenp-C3* gene between the *atms* and *CG14655* genes in *D. vulcana*, which still maintains both paralogs as intact genes.

Except for *D. triauraria* (*montium* subgroup), *D. kikkawai* (*kikkawai* subgroup), and *D. serrata* (*serrata* subgroup) genomes, all other inferences were based on short-read sequencing of 20 genomes from the *montium* group. Recent studies have performed long-read sequencing-based genome assembly of six *montium* group species: *D. auraria* and *D. rufa* (*montium subgroup*), *D. serrata*, *D. bunnanda,* and *D. birchii* (*serrata* subgroup), and *D. jambulina* (*seguyi* subgroup) **(see Supplementary Table 2 for scaffold IDs)** (Kim et al., 2021, 2024; Son et al., 2025). We re-examined these genome assemblies and confirmed the scenario we have outlined (**Figure 4**), with just one exception. In *D. birchii*, we identified four tandemly arranged segments containing *Cenp-C* and neighboring genes. As a result, *D. birchii* encodes two intact *Cenp-C1c* genes, which share 99.7% nucleotide identity, whereas the other two copies are partially duplicated **(Supplementary Figure 6).**

Thus, it appears there might be two *bona fide Cenp-C* paralogs in the *montium* group. The first is a significantly diverged *Cenp-C3* paralog that arose via gene duplication into a new genomic locus, flanked by *atms* and *CG14655* in *D. vulcana* **(Figure 1**; **Figure 4**, step 5). The second is a *Cenp-C1c2* duplicate that arose as part of a 4-gene tandem duplication in *D. birchii* **(Supplementary Figure 6)**. Both these paralogs are very young and likely limited to just a few species, in contrast to the long-term retention of multiple *Cid* paralogs in this group.

## Discussion

Here, we investigate and reject the possibility of an ancient co-retention of *Cid* and *Cenp-C* paralogs in the *montium* group of *Drosophila* species. Although most species in the *montium* group retain only a single functional *Cenp-C* gene, this gene is found in multiple distinct syntenic locations across species, reflecting a history of gene relocation, duplication, and loss rather than stable inheritance at a single locus, as observed in most other *Drosophila* species. Despite its mobility, the phylogeny of various *Cenp-C* genes matches the expected species phylogeny, and the conserved protein motifs essential for centromere function are retained in Cenp-C orthologs across different locations, indicating functional conservation. Thus, for all practical purposes, these *Cenp-C* genes in the *montium* group should be considered functional orthologs. Our findings reveal the unexpected mobility of the *Cenp-C* gene within the *montium group* and suggest that a variety of evolutionary pressures and mechanisms shape the genomic architecture and functional roles of the *Cid* and *Cenp-C* genes across *Drosophila* species.

Our analyses reveal two relatively young *Cenp-C* paralogs. The first of these is the *Cenp-C3* gene in *D. vulcana,* which unambiguously groups with *D. vulcana Cenp-C1c* in our phylogenetic analysis. Given the moderate divergence between these two paralogs, they have been co-retained in *D. vulcana* for a significant period (∼5 MYA) (Yassin, 2018). Since the *seguyi* subgroup, to which *D. vulcana* belongs, is composed of at least 20 species (Yassin, 2018), analysis of more *seguyi* subgroup species might reveal *Cenp-C1c* and *Cenp-C3* co-retention in this entire subgroup. Alternatively, *Cenp-C3* paralog co-retention might be polymorphic even within *D. vulcana* strains. Nevertheless, *Cenp-C3* provides a recent snapshot of the process by which *Cenp-C* might duplicate or relocate to new locations in the *montium* group of species. Although we were unable to identify a *Cenp-C* paralog in *D. birchii* using short-read sequencing, long-read sequencing identified a near-identical paralog (*Cenp-C1c2*) that arose via a tandem duplication. These paralogs are much younger than those in *D. vulcana,* and they are likely polymorphic among different *D. birchii* strains. Short-read sequencing is not ideal for detecting tandem duplication events, and long-read sequencing analyses might reveal more such young duplications in the future (Chakraborty et al., 2021; Kim et al., 2021; Krsticevic et al., 2015; Rogers et al., 2014).

What underlies the unusual mobility of *Cenp-C* genes to new syntenic locations in the *montium* group of species? One potential clue emerges from a closer examination of the most recent duplication of *Cenp-C3* in *D. vulcana*. We found a *Harbinger*-like TIR DNA transposon flanking both *Cenp-C1c* and *Cenp-C3* in *D. vulcana* **(Supplementary Figure 7A)**. It is tempting to speculate that this *Harbinger*-like transposon may have contributed to the *Cenp-C* duplication in *D. vulcana*. TIR DNA transposons are known to cause duplications when the transposase recognizes TIRs from two different TE copies, resulting in duplication of the intervening DNA segment. However, increasing the distance between TIRs reduces the likelihood of interactions required for transposition complex formation, so *Harbinger* elements with excessively long internal sequences are less likely to transpose (Redd et al., 2023). Thus, a large gene such as *Cenp-C*, which spans more than 5 kb, is unlikely to be duplicated by this mechanism. However, proximity to DNA transposons might render *Cenp-C* genes more susceptible to proximal DNA breaks, leading to gene relocation or duplication as a by-product of host DNA repair. Indeed, TEs can mediate the movement of adjacent genomic sequences during transposition events (Feschotte and Pritham 2007).

We further explored the incidence of TEs in and around *Cenp-C* genes. We observed extensive variation in the size of *Cenp-C* genes in the *montium* group, ranging from 5,575 to 10,801 bp, even though *Cenp-C* coding sequences (CDS) varied only from 3,513 to 4,650 bp **(Supplementary Table 3**). Closer examination revealed that TEs correspond to 34.4% (or 3,721 bp) of the *Cenp-C* gene length in *D. watanabei Cenp-C1c*, or to 39.6% (or 3,982 bp) in *D. vulcana Cenp-C3*. Even *Cenp-C* sequences with small sizes have a TE sequence, such as *D. leontia Cenp-C1c* (which has 12.7%, or 706 bp derived from a TE) or *D. pectinifera Cenp-C1b* (8.2%, or 458 bp) **(Supplementary Figure 7B)**. In addition to the full-length *Cenp-C* genes, we also found small fragments of *Cenp-C* in *D. punjabiensis*, *D. vulcana,* and *D. bakoue,* which were flanked by several transposable elements (TEs) (**Supplementary Figure 4)**. All these findings lead to our hypothesis that a high genic TE content may have contributed to the unusual mobility of *Cenp-C* genes in the *montium* group relative to other *Drosophila* species.

Recent studies have highlighted the high content and dynamic activity of transposable elements (TEs) in the *montium* group of *Drosophila* species. For example, research on *D. serrata* showed considerable variation in TE copy number among genotypes and indicated active TE transposition bursts rather than steady accumulation (Tiedeman and Signor 2021). This pattern suggests that TEs are an important and variable component of the genome in *montium* group species. Further comparisons of genic versus heterochromatic TE content might reveal whether the *montium* group has generally high genic TE abundance and whether this has led to the mobility of many other genes, such as *Cenp-C*.

We find the co-retention of *Cid* paralogs, but not *Cenp-C* paralogs, in the *montium* group (*Sophophora* subgenus) intriguing, especially because *Cid* and *Cenp-C* paralogs do appear to have been co-retained in most species of the *Drosophila* subgenus. Our findings imply that only one Cenp-C protein (with only one CenH3 binding motif) interacts with all three Cid variants (Cid1, Cid3, and Cid4) in the *montium* group of *Drosophila* species. Even though both *Cid* and *Cenp-C* are functionally essential for chromosome segregation, we wondered why they differ in their propensity to undergo functional specialization. Distinct evolutionary pressures might drive this difference. Indeed, *Cid1* appears to have evolved under positive selection during the divergence of *D. melanogaster* and *D. simulans* in the *melanogaster* group (Malik & Henikoff, 2001). Similarly, at least two of three *Cid* paralogs appear to have been subject to positive selection in the *montium* group (Kursel et al. 2021). In contrast, we found no evidence of positive selection acting on *Cenp-C* or their chaperone *Cal1* in either *D. melanogaster-D. simulans* or *D. serrata*-*D. bunnanda* from the *montium* group of species **(Supplementary Table 4)**. This rapid evolution of *Cid* might result in a functional trade-off, in which a single *Cid* gene cannot optimally perform all centromeric mitotic and meiotic functions. Such a tradeoff might potentially select for the separation of function via functional specialization of *Cid* gene duplicates, as we observe in *D. virilis* (*Drosophila* subgenus), and is likely the case in the *montium* group as well. There is also similar evidence for CenH3 duplication and sub-functionalization in other animal and plant species (Ishii et al. 2020; Prosée et al. 2020; Evtushenko et al., 2017, 2021; Sanei et al., 2011). However, since *Cenp-C* genes do not undergo positive selection in the *montium* group, there might not be a strong incentive for functional specialization of *Cenp-C* paralogs in *Drosophila* species, leading to a lower probability of co-retention of paralogs that diverge in function or tissue-specific expression. Unlike *Drosophila* species, *Cenp-C* is much more rapidly evolving than *CenH3* in both mammals and plants (Ishii et al., 2020; Talbert et al., 2004). If rapid evolution increases the probability of functional specialization, we might expect to find co-retention and functional specialization among *Cenp-C* paralogs in plant and mammalian species – a possibility that can be explored in future studies.

## Materials and methods

### Identification and annotation of *Cenp-C* genes in the *montium* group

We performed a tBLASTn search on the NCBI database using the sequence of the *D*. *melanogaster* Cenp-C protein as a query to obtain the corresponding mRNAs in *Drosophila* species from three species of the *montium* group, for which mRNA or transcriptomic data were available (*D. serrata*, *D. kikkawai,* and *D. triauraria*) **(Supplementary Table 2)**. The retrieved mRNAs were then used to reconstruct Cenp-C protein sequences, which were used to identify *Cenp-C* gene(s) in the genomes of all species in the *montium* group, using the Cenp-C protein sequence from the most closely related species as a query. We used the Augustus program (Stanke & Morgenstern, 2005) to predict the *Cenp-C* coding sequences (CDS) from each gene hit retrieved from tBLASTn searches.

To access the chromosome location of all *Cenp-C* loci, we used the available chromosome-length scaffolds from *D. triauraria* (*montium* subgroup) as a reference (Torosin et al., 2020). All gene names mentioned in this work are consistent with the annotations of *D. melanogaster* available on FlyBase 2.0 (Thurmond et al., 2019). Transposable elements flanking or within Cenp-C were identified using the CENSOR tool available at the Repbase database (Kohany et al., 2006).

### Phylogenetic trees

The *Cenp-C* coding sequences of the *Drosophila* species were aligned using the MUSCLE algorithm implemented in Geneious Prime (https://www.geneious.com), followed by manual refinement. Since *Cenp-C* mRNA or transcriptomic data are unavailable for most *montium* group species, exon sequences absent or duplicated in one or more species (see **Supplementary Figure 4**) were excluded from the phylogenetic analysis. The phylogenetic trees were inferred using the maximum likelihood method and the nucleotide substitution model GTR+G+I, which is the best substitution model for our data. These analyses were implemented on MEGAX software (Kumar et al., 2018) using 1000 bootstrap replicates.

### Analysis of the Cenp-C protein motifs

Seven previously identified motifs from *D. melanogaster* Cenp-C (Heeger et al., 2005) were used to search the Cenp-C protein sequence in species from the *montium* group. We aligned the amino acid sequences of Cenp-C from *D. melanogaster* and nine other species belonging to the *melanogaster* group (*D. simulans*, *D. sechellia*, *D. mauritiana, D. yakuba*, *D. erecta*, *D. eugracilis, D. elegans*, *D. ficusphila*, and *D. takahashii*) using the MUSCLE algorithm implemented in Geneious Prime. The *Cenp-C* mRNA of the species used in this analysis is available on the NCBI database **(Supplementary Table 2)**.

We then identified and extracted the sequence regions corresponding to each *D*. *melanogaster* Cenp-C motif on the alignment. We selected conserved motifs (with at least 60% identity) compared with the same motif in other species within the *Drosophila melanogaster* group. The motif discovery algorithm *MEME* (Bailey et al., 2015) was used to generate position-specific scoring matrices for each motif, which describe the probability of each amino acid at each position in the pattern. Sequences containing indels were excluded from the analysis, as MEME does not account for indels. Each matrix was used for motif scanning in Cenp-C from the *montium* group species using the *MAST* algorithm (Bailey & Gribskov, 1998). Only motifs with *p*-value < 10^-6^ were considered. The returned sequence matches, and the sequences used in the first search were used together to produce new matrices for a second round of scanning. Again, only sequences with *p*-value < 10^-6^ were considered.

### Positive selection analysis

We applied McDonald–Kreitman tests to compare within-species polymorphism and between-species divergence (McDonald and Kreitman, 1991) to look for positive selection in two individual lineages, *D. melanogaster* and *D. serrata*. We used *D. simulans* as the closely related outgroup species for the *D. melanogaster* analysis, and *D. bunnanda* for the *D. serrata* analysis, except for *Cid3* (which degenerated in *D. bunnanda)*, for which we used *D. birchii* as the outgroup. We extracted population data from public datasets of >1000 *D. melanogaster* strains (Hervas et al., 2017; Lack et al., 2016) and 111 *D. serrata* strains (Reddiex et al., 2018). We conducted unpolarized McDonald–Kreitman tests using R scripts (https://github.com/jayoung/MKtests_JY).

## Data availability

All data supporting the findings of this study are available within the article and its supplementary information files.

