## Supplementary Material for "Retention of a single *Cenp-C* gene in different syntenic locations in the *montium* group of *Drosophila* species"

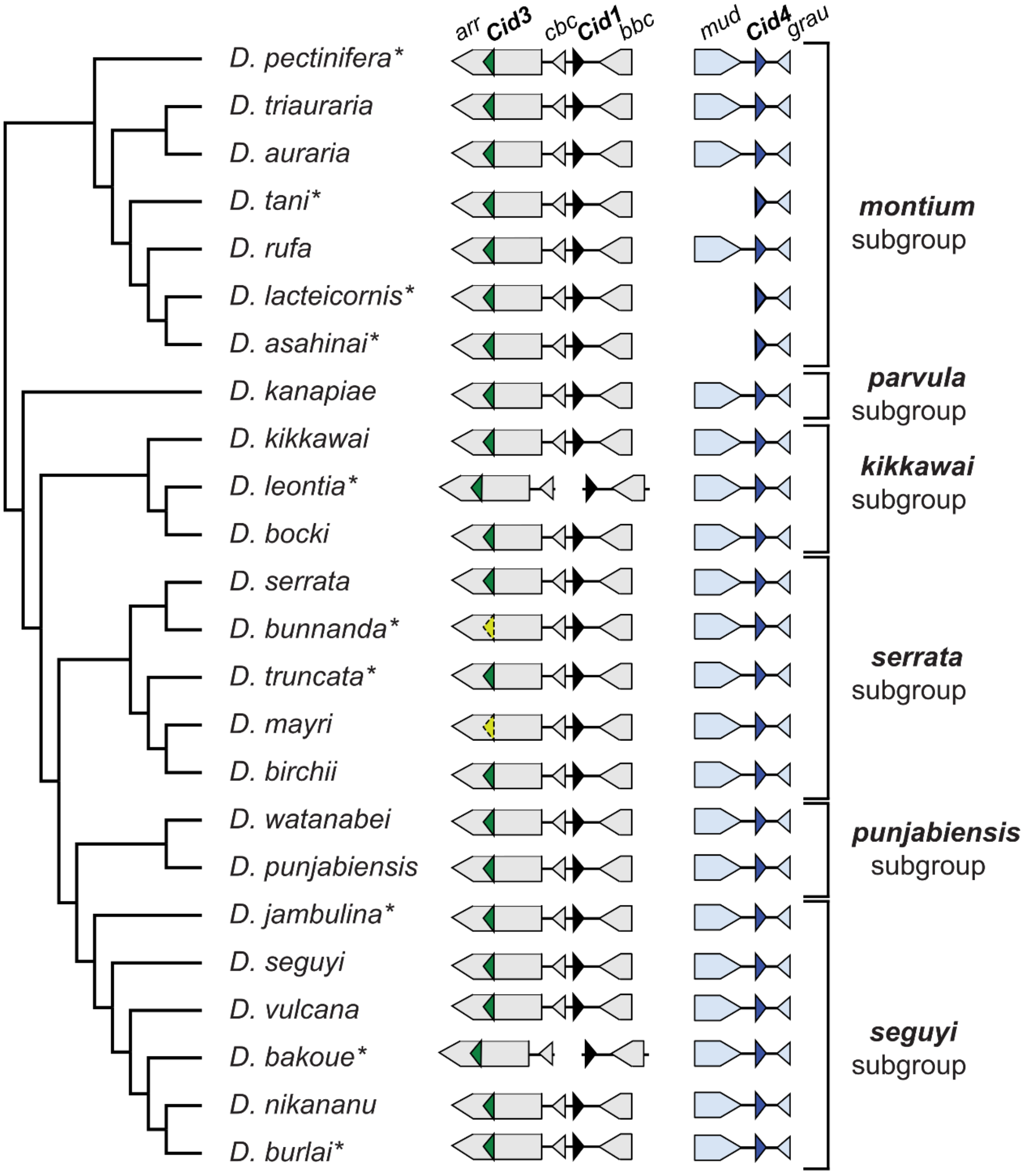


#### Figure S1

Phylogeny of the 24 species from the *montium* group (adapted from Conner et al., 2021) analyzed in the present work, with a schematic representation of their *Cid* paralogs. The *Cid* paralogs and their shared syntenic loci (represented by the flanking genes) are shown for all species, 14 of which were previously reported (Kursel & Malik, 2017). The additional ten species analyzed here are indicated with an asterisk. Contig breaks reflect limitations associated with short-read genome assemblies in some species.

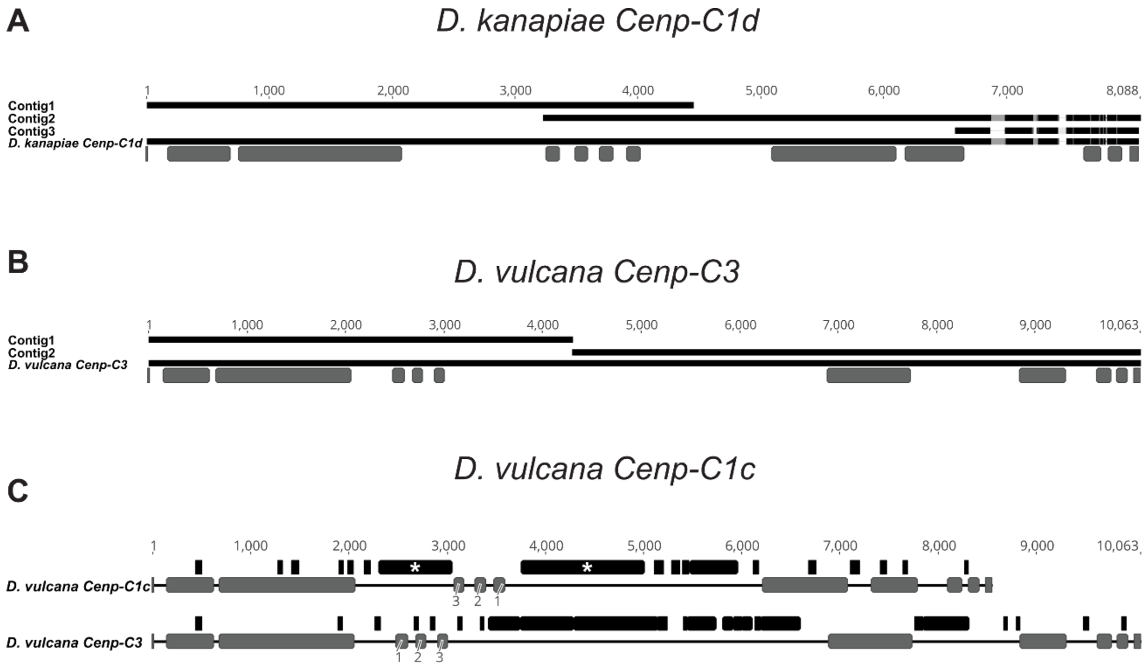


### Figure S2

**Predicted *Cenp-C* gene sequences for *D. kanapiae* and *D. vulcana*. (A)** Alignment of the three *Cenp-C* gene fragments in *D. kanapiae* (contigs 1, 2, and 3) with the predicted *Cenp-C1d* sequence. Identical regions (black), divergent regions (gray), and gaps (white) are shown in the alignment. Gray boxes at the bottom represent exons. **(B)** Alignment of the two *Cenp-C* gene fragments in *D. vulcana* (contigs 1 and 2) with the predicted *Cenp-C3* sequence. **(C)** The *D. vulcana Cenp-1c* gene includes three exons (numbered 1-3) that appear to be inverted within a predicted intron sequence of the *Cenp-C1c* gene, flanked by transposable elements (*Helitron*-like elements, represented as a black bar with an asterisk). Black bars indicate sequences with similarity to TEs according to the RepBase database.

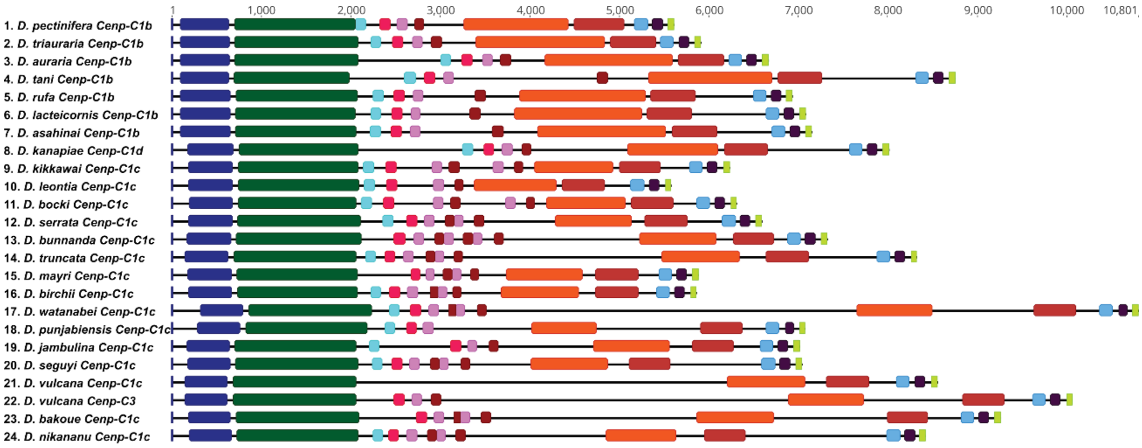


### Figure S3

***Cenp-C* genes from the *montium* group.** Schematic representation of all *Cenp-C* genes, with exonic sequences shown as colored bars. Bars with the same color correspond to the equivalent sequence across *Cenp-C* genes. Exonic sequences located between the green and the orange bars in each gene represent regions that were excluded from the phylogenetic analysis because they could not be found in all species.

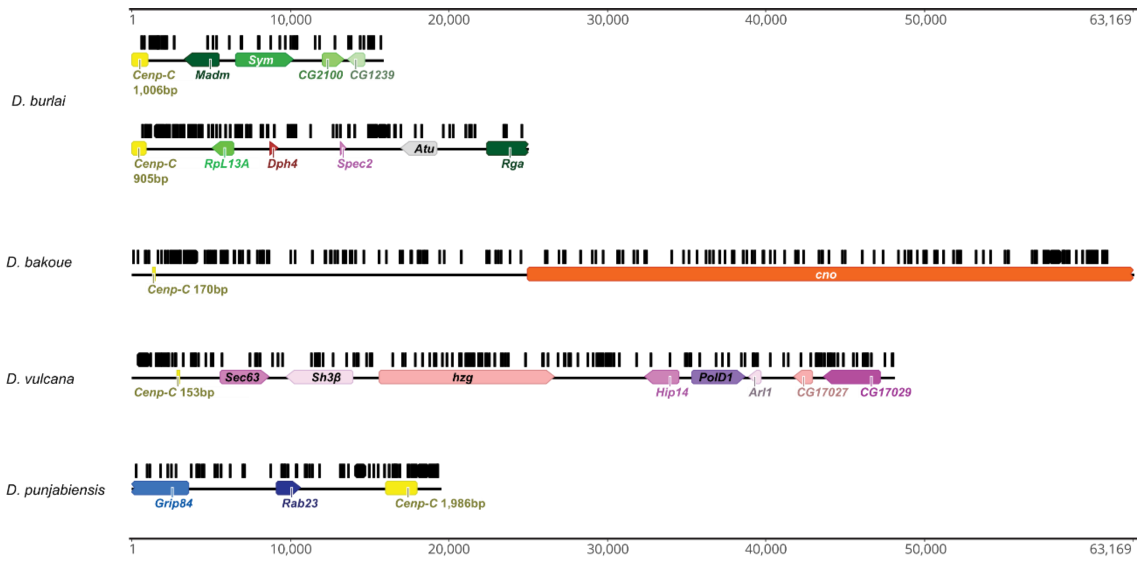


### Figure S4

**Additional *Cenp-C* fragments are found in distinct genomic regions across four species of the *montium* group.** Genes and *Cenp-C* fragments are represented by colored bars, and intergenic sequences by black lines. Black bars indicate sequences with similarity to transposable elements, according to the RepBase database. Due to contig breaks, the genes *Rga*, *cno*, and *Grip84* are incomplete.

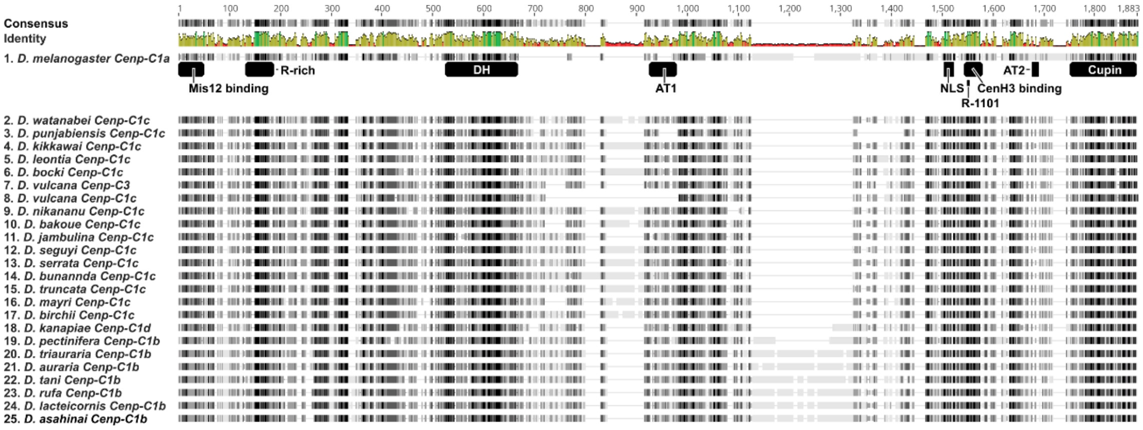


### Figure S5

**Alignment of Cenp-C amino acid sequences from species of the *montium* group showing the positions of the protein motifs identified in this study.** The black bars represent, from left to right: Mis12 binding, R-rich, DH, AT1, NLS, CenH3 binding, AT2, and Cupin. The small bar below the CenH3 binding motif indicates the R-1101 amino acid, which is present in all Cenp-C genes from the *montium* group. Motifs AT1 and AT2 were not detected in species from the *montium* group. MEME was used to identify the Mis12 binding motif, while MAST analyses detected all the other motifs.

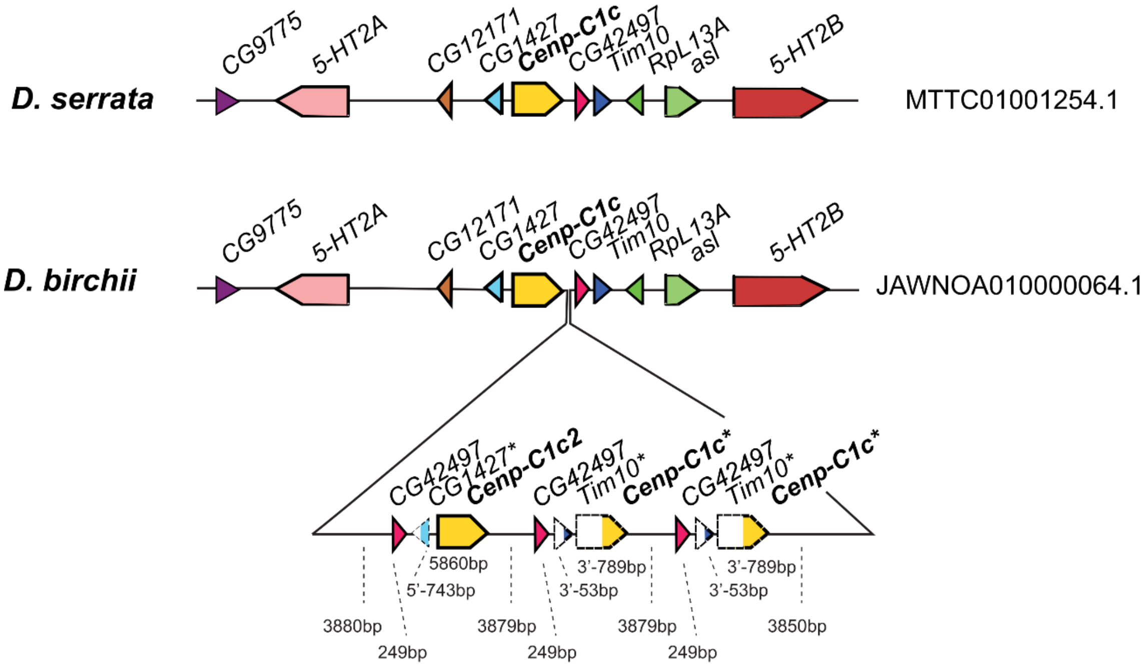


### Figure S6

**Tandem duplications in the *Cenp-C1c* locus of *D. birchii* revealed by long-read assembly analysis.** For comparison, the corresponding locus is shown for *D. serrata*, which also belongs to the *serrata* subgroup and for which a long-read assembly was also analyzed. The tandem duplication involved both genes and intergenic sequences. Some genes were wholly duplicated, while others were partially duplicated (asterisk). For example, this locus now encodes four intact copies of *CG42497*, but only two intact copies of *Cenp-C1c*, and no additional copies of flanking genes *CG1427, Tim10,* and *RpL13A.*

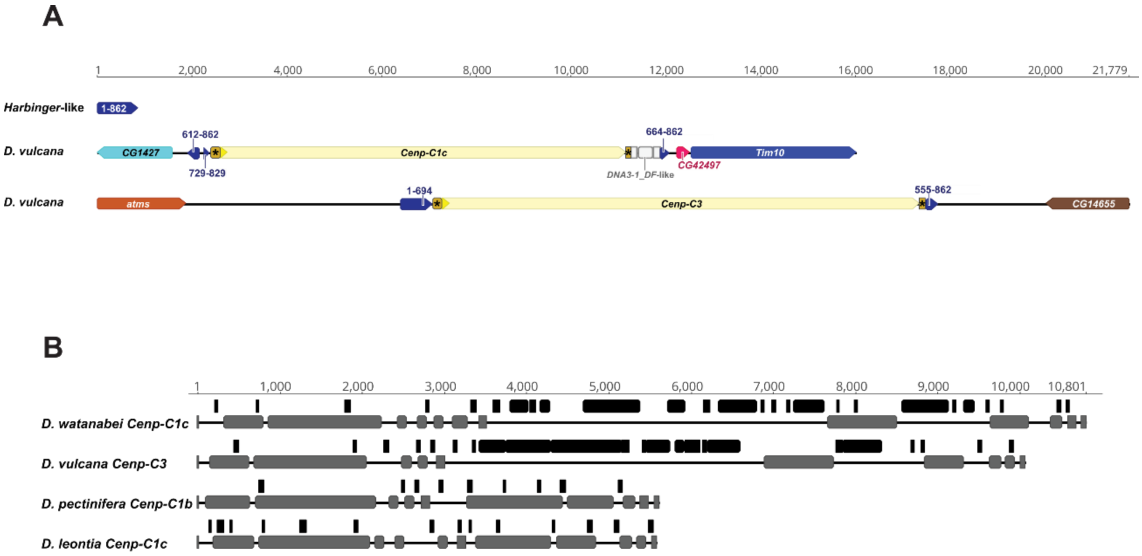


### Figure S7

**High density of TEs within the *Cenp-C* sequence and locus**. **(A)** A repetitive sequence of approximately 862 bp is found among species of the *D. seguyi* subgroup. This sequence shows similarity to *Harbinger* elements (Repbase). Therefore, we annotate it as a *Harbinger*-like element. Segments of this Harbinger-like element flank *Cenp-C1c* and *Cenp-C3* on both sides in *D. vulcana*. The yellow bar marked with an asterisk indicates the 5′ and 3′ sequences (likely corresponding to the 5′ and 3′ UTRs) that exhibit similarity between *Cenp-C1c* and *Cenp-C3* in *D. vulcana*. The sequence between the *Cenp-C1c* 3′ UTR and the *Harbinger*-like element corresponds to a distinct DNA transposon (*DNA3-1_DF*-like element). **(B)** Enrichment of transposable elements within introns of *Cenp-C*. Schematic representation of *Cenp-C* genes from *D. watanabei* (*Cenp-C1c*), *D. vulcana* (*Cenp-C3*), *D. pectinifera* (*Cenp-C1b*), and *D. leontia* (*Cenp-C1c*). The gray bars represent exons, and the black bars represent sequences with similarity to TEs**.**

### Supplementary Tables

#### **Supplementary Table 1.** Genbank Database IDs

| Species | *Muller element E* | Muller element C |
|---|---|---|
| ***D. jambulina*** | VNJX01008392.1 | VNJX01003011.1 |
| ***D. watanabei*** | VNJS01015544.1 | VNJS01016027.1 |
| ***D. serrata*** | MTTC01001254.1 | MTTC01000143.1 |
| ***D. kikkawai*** | JARPSD010000005.1 | JARPSD010000003.1 |
| ***D. kanapiae*** | VNJM01000172.1 | VNJM01002302.1 |
| ***D. triauraria*** | JABJVT010000005.1 | JABJVT010000003.1 |
| ***D. melanogaster*** | NT_033777.3 | NT_033778.4 |
| ***D. ananassae*** | JACRYV020000138.1 | JACRYV020000136.1 |
| ***D. virilis*** | JAUDTK010000001.1 | JAUDTK010000004.1 |
| ***S. lebanonensis*** | QMEN02000020.1 | QMEN02000001.1 |

**Supplementary Table 2.** Genbank Database IDs

| Gene | Scaffold or genomic SRA | mRNA or transcriptomic SRA |
|---|---|---|
| ***D. pectinifera Cenp-C1b*** | VNKC01003598.1 | Unavailable |
| ***D. triauraria Cenp-C1b*** | JABJVT010000005.1 | SRR11780982 |
| ***D. auraria Cenp-C1b*** | VNJW01009994.1 and **JBNRRN010000002.1** | Unavailable |
| ***D. tani Cenp-C1b*** | VNJO01011109.1 | Unavailable |
| ***D. rufa Cenp-C1b*** | VNKH01000388.1 and **JAECXS010000001.1/ JAECXS010000038.1** | Unavailable |
| ***D. lateicornis Cenp-C1b*** | VNKF01009708.1 | Unavailable |
| ***D. asahinai Cenp-C1b*** | VNJZ01004887.1 | Unavailable |
| ***D. kanapiae Cenp-C1d*** | VNJM01000860.1/ VNJM01001987.1 | Unavailable |
| ***D. kikkawai Cenp-C1c*** | AFFH02006098.1 | XM_017182092.1 |
| ***D. leontia Cenp-C1 c*** | VNKB01009439.1 | Unavailable |
| ***D. bocki Cenp-C1 c*** | VNJY01005756.1 | Unavailable |
| ***D. serrata Cenp-C1 c*** | MTTC01001254.1 and JAWNOG010000382.1/JAWNOG010000364.1/JAWNOG010000228.1 | XM_020957133.1 |
| ***D. bunnanda Cenp-C1 c*** | VNKE01001283.1 and **JAWNOB010000030.1/JAWNOB010000130.1/JAWNOB010000100.1** | Unavailable |
| ***D. truncata Cenp-C1 c*** | VNJQ01004447.1 | Unavailable |
| ***D. mayri Cenp-C1 c*** | VNJN01008118.1 | Unavailable |
| ***D. birchii Cenp-C1 c*** | VNKA01006487.1 and **JAWNOA010000064.1** | Unavailable |
| ***D. watanabei Cenp-C1 c*** | VNJS01015544.1 | Unavailable |
| ***D. punjabiensis Cenp-C1 c*** | VNJR01008397.1 | Unavailable |
| ***D. jambulina Cenp-C1 c*** | VNJX01008392.1 and **JAECXH010000012.1** | Unavailable |
| ***D. seguyi Cenp-C1 c*** | VNJU01009249.1 | Unavailable |
| ***D. vulcana Cenp-C1 c*** | VNJP01006877.1 | Unavailable |
| ***D. vulcana Cenp-C3*** | VNJP01005529.1/ VNJP01004469.1 | Unavailable |
| ***D. bakoue Cenp-C1c*** | VNJL01004438.1 | Unavailable |
| ***D. nikananu Cenp-C1c*** | VNJV01009586.1 | Unavailable |
| ***D. melanogaster Cenp-C1a*** | NT_033777.3 | NM_169228.3 |
| ***D. simulnas Cenp-C1a*** | NIFY01000004.1 | XM_016174981.1 |
| ***D. sechellia Cenp-C1a*** | NIFZ01000004.1 | XM_032720766.1 |
| ***D. yakuba Cenp-C1a*** | JAEDAC010000005.1 | XM_002096785.2 |
| ***D. erecta Cenp-C1a*** | QMER02000001.1 | XM_026983729.1 |
| ***D. eugracilis Cenp-C1a*** | AFPQ02005741.1 | SRR346729 |
| ***D. elegans Cenp-C1a*** | WVIB01000005.1 | XM_017268702.1 |
| ***D. ficusphila Cenp-C1a*** | AFFG02008693.1 | XM_017203168.1 |
| ***D. takahashii Cenp-C1a*** | XM_017148281.1 | AFFI02007576.1 |
| ***D. mauritiana Cenp-C1a*** | XM_033307073.1 | NIGA01000004.1 |
| ***D. ananassae Cenp-C1a*** | JACRYV010000154.1 | XM_001955206.3 |
| ***D. persimilis Cenp-C1a*** | QMET02000001.1 | XM_026986294.1 |
| ***D. pseudoobscura Cenp-C1a*** | WVEN01000001.1 | XM_015182067.2 |
| ***D. willistoni Cenp-C1a*** | AAQB01009414.1 | XM_023180684.1 |
| ***D. mojavensis Cenp-C1a*** | CH933806.1 | SRR6425997 |
| ***D. mojavensis Cenp-C2*** | CH933806.1 | SRR6425997 |
| ***D. arizonae Cenp-C1a*** | SRR2070760 | SRR2509638 |
| ***D. arizonae Cenp-C2*** | SRR2070760 | SRR2509638 |
| ***D. navojoa Cenp-C1a*** | LSRL02000055.1 | Unavailable |
| ***D. navojoa Cenp-C2*** | LSRL02000228.1 | XM_018113591.2 |
| ***D. buzzatii Cenp-C1a*** | https://dbuz.uab.cat/blast.php *D. buzzatii* Freeze 1 Scaffolds | SRR5145562/ SRR5145563 |

| | | |
|---|---|---|
| ***D. buzzatii Cenp-C2*** | https://dbuz.uab.cat/blast.php *D. buzzatii* Freeze 1 Scaffolds | SRR5145562/ SRR5145563 |
| ***D. seriema Cenp-C1a*** | ERR1976657 | Unavailable |
| ***D. seriema Cenp-C2*** | ERR1976657 | Unavailable |
| ***D. virilis Cenp-C1a*** | QMEO02000199.1 | XM_002056576.3 |
| ***D. virilis Cenp-C2*** | QMEO02000199.1 | XM_002056451.3 |
| ***D. americana Cenp-C1a*** | UEJX01001328.1 | SRR5279019 |
| ***D. americana Cenp-C2*** | UEJX01001683.1 | SRR5279019 |
| ***D. grimshawi Cenp-C1a*** | AAPT01020190.1 | XM_001994049.2 |
| ***D. grimshawi Cenp-C2*** | AAPT01019320.1 | XM_001989754.3 |
| ***D. busckii Cenp-C1a*** | DULD01000002.1 | XM_017991761.1 |
| ***D. busckii Cenp-C2*** | DULD01000002.1 | XM_017992885.2 |
| ***Phortica variegata Cenp-C1a*** | JXPM01003917.1 | SRR1738675 |

NOTE: The long-read sequencing-based genome assemblies for six species are shown in bold.

**Supplementary Table 3.** General characteristics of the *Cenp-C* genes in species from the *montium* group.

| Subgroups | Gene | nº of exons | gene length (bp) | CDS length (bp) |
|---|---|---|---|---|
| *montium* | *D. pectinifera Cenp-C1b* | 10 | 5,610 | 4,386 |
| | *D. triauraria Cenp-C1b* | 11 | 5,907 | 4,653 |
| | *D. auraria Cenp-C1b* | 11 | 6,667 | 4,635 |
| | *D. tani Cenp-C1b* | 11 | 8,750 | 4,488 |
| | *D. rufa Cenp-C1b* | 11 | 6,933 | 4,599 |
| | *D. lacteicornis Cenp-C1b* | 11 | 7,078 | 4,590 |
| | *D. asahinai Cenp-C1b* | 11 | 7,147 | 4,605 |
| *parvula* | *D. kanapiae Cenp-C1d* | 11 | 8,013 | 4,089 |
| *kikkawai* | *D. kikkawai Cenp-C1c* | 13 | 6,229 | 4,152 |
| | *D. leontia Cenp-C1c* | 11 | 5,575 | 3,981 |
| | *D. bocki Cenp-C1c* | 13 | 6,310 | 4,125 |
| *serrata* | *D. serrata Cenp-C1c* | 12 | 6,591 | 4,152 |
| | *D. bunnanda Cenp-C1c* | 12 | 7,322 | 4,248 |
| | *D. truncata Cenp-C1c* | 12 | 8,324 | 4,131 |
| | *D. mayri Cenp-C1c* | 11 | 5,885 | 3,957 |
| | *D. birchii Cenp-C1c* | 12 | 5,861 | 4,104 |
| *punjabiensis* | *D. watanabei Cenp-C1c* | 12 | 10,801 | 4,068 |
| | *D. punjabiensis Cenp-C1c* | 10 | 7,076 | 3,657 |
| *seguyi* | *D. jambulina Cenp-C1c* | 11 | 7,015 | 3,867 |
| | *D. seguyi Cenp-C1c* | 12 | 7,039 | 4,119 |
| | *D. vulcana Cenp-C1c* | 7* | 8,552 | 3,516 |
| | *D. vulcana Cenp-C3* | 10 | 10,063 | 3,801 |
| | *D. bakoue Cenp-C1c* | 11 | 9,264 | 3,981 |
| | *D. nikananu Cenp-C1c* | 12 | 8,420 | 4,026 |

NOTE: The *Cenp-C1c* from *D. vulcana* has 7 exons according to Augustus algorithm. Our analysis further revealed that *Cenp-C1c* has 10 exons (see **Supplementary Figure 2C** for more details).

**Supplementary Table 4:** McDonald-Kreitman tests for positive selection

| Species | Gene | FET p-value | Dn | Ds | Pn | Ps | NI | α |
|---|---|---|---|---|---|---|---|---|
| ***D. melanogaster* vs *D. simulans*** (***melanogaster*** **group**) | ***Cid1*** $^{NR}$ | **0,015** | 22 | 19 | 1 | 9 | 0,096 | 0,904 |
| | ***CenpC*** $^{NR}$ | 0,815 | 121 | 101 | 11 | 8 | 1,148 | -0,148 |
| | ***Cal1*** $^{NR}$ | 0,757 | 80 | 77 | 27 | 30 | 0,866 | 0,134 |
| ***D. serrata*** **vs** ***D. bunnanda*** (***montium*** **group**) | ***Cid1*** | **0,010** | 39 | 33 | 8 | 23 | 0,294 | 0,706 |
| | ***Cid1*** $^{NR}$ | 0,088 | 39 | 34 | 2 | 8 | 0,218 | 0,782 |
| | ***Cid3*** | 0,201 | 54 | 52 | 16 | 26 | 0,593 | 0,407 |
| | ***Cid3*** $^{NR}$ | 0,144 | 55 | 54 | 6 | 13 | 0,453 | 0,547 |
| | ***Cid4*** | 0,186 | 27 | 27 | 22 | 38 | 0,579 | 0,421 |
| | ***Cid4*** $^{NR}$ | **0,010** | 28 | 29 | 2 | 15 | 0,138 | 0,862 |
| | ***CenpC*** | 0,135 | 160 | 175 | 67 | 52 | 1,409 | -0,409 |
| | ***CenpC*** $^{NR}$ | 0,155 | 162 | 177 | 12 | 23 | 0,570 | 0,430 |
| | ***Cal1*** | 0,811 | 131 | 137 | 190 | 189 | 1,051 | -0,051 |
| | ***Cal1*** $^{NR}$ | 0,408 | 142 | 146 | 62 | 77 | 0,828 | 0,172 |

NOTE. We carried out unpolarized McDonald-Kreitman tests (McDonald and Kreitman, 1991) of the *Cid, Cenp-C,* and *Cal1* genes across ~1000 *D. melanogaster* strains, using *D. simulans* as an outgroup. Since many sequenced strains were available, we ignored polymorphisms at less than 5% frequency, which are less likely to have been tested by natural selection (Fay et al. 2002); these genes are indicated with an *NR* (no-rare) superscript. In the *melanogaster-simulans* comparison, only *Cid* shows evidence of positive selection ($p<0.05$; bold), consistent with a previous study (Malik & Henikoff, 2001). We then compared the *Cid1, Cid3, Cid4, CenpC,* and *Cal1* genes from ~110 *D. serrata* sequences to those from *D. bunnanda*. However, since *D. bunnanda Cid3* is a pseudogene, we used *D. birchii Cid3* as an outgroup. We analyzed data for all polymorphisms or using comparisons that ignore rare polymorphisms (NR superscript). Our data finds evidence for positive selection in *Cid1* and *Cid4* in one of the two comparisons ($p<0.05$; bold), but no evidence for positive selection for *Cid3*, *CenpC*, or *Cal1* in either comparison.


Fay, J. C., Wyckoff, G. J., & Wu, C. I. (2002). Testing the neutral theory of molecular evolution with genomic data from Drosophila. *Nature*, *415*(6875), 1024–1026. https://doi.org/10.1038/4151024a

Malik, H. S., & Henikoff, S. (2001). Adaptive evolution of Cid, a centromere-specific histone in Drosophila. *Genetics*, *157*(3), 1293–1298. https://doi.org/10.1093/genetics/157.3.1293

McDonald, J. H., & Kreitman, M. (1991). Adaptive protein evolution at the Adh locus in Drosophila. *Nature*, *351*(6328), 652–654. https://doi.org/10.1038/351652a0
